# Leveraging genomics and temporal high-throughput phenotyping to enhance association mapping and yield prediction in sesame

**DOI:** 10.1101/2024.02.01.578346

**Authors:** Idan Sabag, Ye Bi, Maitreya Mohan Sahoo, Ittai Herrmann, Gota Morota, Zvi Peleg

## Abstract

Sesame (*Sesamum indicum*) is an important oilseed crop with rising demand due to its high oil quality. To meet these future demands, there is an urgent need to develop and integrate new breeding strategies. While genomic resources have advanced genetic research in sesame, implementation of high-throughput phenotyping and genetic analysis of longitudinal traits remains limited. Here, we combined high-throughput phenotyping and random regression models to investigate the dynamics of plant height, leaf area index, and five spectral vegetation indices throughout the sesame growing seasons in a diversity panel. Modeling the temporal phenotypic and additive genetic trajectories revealed distinct patterns corresponding to the sesame growth cycle. We also conducted longitudinal genomic prediction and association mapping of plant height using various models and cross-validation schemes. Moderate prediction accuracy was obtained when predicting new genotypes at each time point, and moderate to high values were obtained when forecasting future phenotypes. Association mapping revealed three genomic regions in linkage groups 6, 8, and 11 conferring trait variation over time and growth rate. Furthermore, we leveraged correlations between the temporal trait and seed-yield and applied multi-trait genomic prediction. We obtained an improvement over single-trait analysis, especially when phenotypes from earlier time points were used, highlighting the potential of using a high-throughput phenotyping platform as a selection tool. Our results shed light on the genetic control of longitudinal traits in sesame and underscore the potential of high-throughput phenotyping to detect a wide range of traits and genotypes that can inform sesame breeding efforts to enhance yield.

## 1. INTRODUCTION

Sesame (*Sesamum indicum*, 2n = 2x = 26) is an ancient oilseed crop whose seeds are rich in high-quality oil, protein, minerals, and antioxidants. With an annual production of 6.8 million tons (2022; https://www.fao.org/faostat/en/#data/QCL) and increasing demands, major breeding efforts should be made to meet these requirements. In recent years, genetic and scientific research in sesame has been greatly improved through the integration of genomic resources and other omics technologies (Weldemichael and Gebremedhn, 2023). Moreover, quantitative trait loci (QTL) conferring a wide range of agronomically important traits have been identified in sesame, including phenological, morphological and yield components (Teboul et al., 2020; Sabag et al., 2021), biotic and abiotic stresses (Dossa et al., 2019; Asekova et al., 2021), and grain quality traits (Teboul et al., 2020; Cui et al., 2021; Dossou et al., 2023). Nevertheless, there is limited information about the genetic components that control the dynamics of longitudinal traits, such as canopy cover, plant height (PH), and other developmental traits. Because sesame is an indeterminate crop-plant, uncovering the genetic architecture of growth patterns is essential to maximize seed-yield.

High-throughput phenotyping (HTP) has revolutionized the process of crop-plant breeding by providing tools for rapid and precise assessment and selection of superior genotypes for a wide range of functional traits (Yang et al., 2020). This approach leverages advanced technologies, such as remote and proximal sensing, imaging, and sensor-based data collection, to gather extensive data on functional traits (e.g., morphological, anatomical, physiological, and biochemical phenotypes). By automating data acquisition and analysis, plant breeders can evaluate large populations more efficiently than with traditional methods (Cabrera-Bosquet et al., 2012). In combination with genomic information, HTP can accelerate the identification of desirable traits, facilitating the selection of superior climate-resilience varieties with enhanced yield and nutritional quality (Mir et al., 2019; Shorinola et al., 2024). In addition, HTP allows the collection of time-series measurements that track the crop-plant development throughout the growing season to exploit the genetic architecture at different stages of crop development. Furthermore, this type of information can be used as a secondary trait for indirect selection and prediction of the target trait by exploiting the correlation between the traits (Rutkoski et al., 2016; Sun et al., 2017). In addition, spectral vegetation indices (SVI) can provide information about plant status and development (Tayade et al., 2022) and previous studies suggested that SVI calculated from HTP could increase the efficiency of indirect selection for seed yield and can be integrated into sesame breeding programs (Petsoulas et al., 2022).

Random regression (RR) models, first introduced in animal breeding (Schaeffer, 2004), use covariance functions, such as orthogonal polynomials and splines, to model the trait trajectory over different time points with few parameters to estimate in a computationally efficient time. This approach considers the covariance between time points, and genomic estimated breeding values (GEBV) can be calculated for each individual at each time point (Mrode, 2014). The integration of HTP data into this framework allowed the investigation of the genetic architecture of shoot biomass in rice (*Oryza sativa*) (Campbell et al., 2018, 2019; Baba et al., 2020), soybean (*Glycine max* ) (Freitas Moreira et al., 2021), and the examination of the response to environmental cues (Sakurai et al., 2023; Rebollo et al., 2023). Here, we combined HTP and genomic data to dissect the genetic composition of longitudinal traits in a sesame diversity panel and utilize this information for genome-wide association mapping (GWAS) and genomic prediction of seed yield under field conditions. The specific objectives of the current study were to: (***i*** ) model the phenotypic and genetic dynamics of morphological traits and spectral vegetation indices along the sesame growing season, (***ii*** ) perform time-series mapping and genomic prediction plant height, and (***iii*** ) integrate genomic and temporal HTP data to improve seed-yield prediction. Our findings indicate that uncovering the genetic composition of longitudinal traits could prove beneficial in advancing sesame research and breeding efforts.

## 2. MATERIALS AND METHODS

### 2.1 Plant material and field experiment conditions

A sesame diversity panel (SCHUJI) containing 185 genotypes from a wide range of geographical origins was used for the current study. The SCHUJI panel was previously genotyped using genotyping-by-sequencing and subjected to phenotypic evaluation, GWAS, and genomic prediction in two field experiments (Sabag et al., 2021, 2023). All measurements described below are attributed to the 2020 growing season, located at the experimental farm of the Hebrew University of Jerusalem (Rehovot, Israel). The experimental design was a complete randomized block with five replicates, where each replicate was a 2.5 (L) and 0.8 (W) meter plot with three rows and a plant density of 6 plants per meter. After quality control, a total of 20,340 single nucleotide polymorphism (SNPs) markers remained for genomic analyses. This process included the removal of tightly linked markers (*r*^2^ *≥* 0.99), minor allele frequencies less than 0.05, and heterozygosity rates greater than 0.2. A full description of the panel development and genotyping procedure can be found at Sabag et al. (2021).

### 2.2 High-throughput phenotyping data collection and analysis

High-throughput phenotyping data collection was conducted at reproductive stages in the sesame growth cycle. To estimate plant development and phenotypic trajectory throughout the growing season, we collected HTP data at six time points (TP): 45, 57, 65, 71, 80, and 87 days after sowing (DAS). Data were collected using both UAV-borne RGB and hyperspectral cameras, as follows.

#### 2.2.1 Plant height estimation

The PH trajectory during the growing season was collected in a few steps. First, images were acquired using a Mavic Mini unmanned aerial vehicle (UAV; DJI, Shenzhen, Guangdong, China) equipped with a 12.7-megapixel RGB camera with bands at 620–750 nm (R, red), 495–570 nm (G, green), and 450–495 nm (B, blue). The UAV images were collected on clear and sunny days between 11:00 and 14:00 local time (solar altitude angle *>* 45^*°*^). The flights were performed 20m above ground level, with 80% image overlap along flight corridors. In the next step, orthomosaic photos and a digital surface model (DSM) were generated using Pix4DMapperPro desktop software (Pix4D SA, Switzerland) with the structure-from-motion algorithm. A canopy height model was completed as described in previous studies (Aharon et al., 2020; Avneri et al., 2023) according to Wallace et al. (2017). To distinguish between vegetation and soil, we calculated the Excessive Green vegetation index (Woebbecke et al., 1995) for the orthomosaic photo. Then we applied the Outsu threshold (Otsu, 1979) in ArcGIS Pro (ESRI, Redlands, CA, USA), which resulted in a binary image between vegetation and soil. To eliminate the soil pixels from the image, we used the Set Null function in the Spatial analysis package in ArcGIS Pro and projected the results on the DSM. A shapefile polygon was constructed, and summary statistics were calculated using the zonal statistics as a table function for each plot in ArcGIS Pro. To validate the results of the image analysis, we manually measured plant height in 200 plots (5 plants each) and compared them to the 95 quantiles (PCT95) values from the summary statistics. The *R*^2^ between these two values was 0.92 (Figure S1). We used PCT95 to represent PH values for all dates for all plots. In addition to the RGB imaging, we monitored the field with a hyperspectral camera.

#### 2.2.2 Hyperspectral image data collection and analysis

UAV-borne hyperspectral imagery was acquired in six flights (one for each time point) of the campaign throughout the growing season using a Pika-L hyperspectral push broom scanner camera (Resonon, Bozeman, MT, USA) mounted on a Hexacopter, Matrice 600 Pro (DJI, Shenzen, China). The camera recorded raw data in the visible and near-infrared regions (380-1020 nm). Data preprocessing included georectification, radiometric correction, and conversion to reflectance using Spectronon Pro software (Resonon, Bozeman, MT, USA). ArcGIS Pro was used to generate polygons for each plot. To make sure that only reflectance related to the plot is included in the polygon, a buffer of 10 cm was used. to minimize the soil effect on the reflectance data, the plot-level reflectance was calculated as the mean of all pixels contained within a polygon after masking all pixels with Normalized Difference Vegetation Index (NDVI) values below 0.85. We filtered the beginning and the end of the spectra and smoothed the values with the Savitzky-Golay smoothing filter with second-order polynomial and default window size using the signal package (signal developers, 2014) in R statistical software (R Core Team, 2022). Wavelengths (WL) between 415 and 930 nm (total of 244) were retained for further analysis.

#### 2.2.3 Leaf area index estimation

We estimated the leaf area index (LAI) for individual plots by combining measured LAI values with their hyperspectral reflectance of the canopy. LAI was manually measured with an AccuPAR LP-80 ceptometer (Decagon Devices, Washington, USA) according to the user manual and based on the photosynthetically active radiation that was intercepted above and below the canopy (Herrmann et al., 2020). In brief, we used a dataset of 120 samples of averaged plot canopy reflectance acquired in two independent time points (45 and 57 DAS) and their measured LAI values. A regression model was developed by training their spectral features using the genetic algorithm-based partial least squares regression. More information regarding LAI estimation can be found in Supplementary Method S1.

#### 2.2.4 High-throughput phenotyping data analysis

Best linear unbiased estimates (BLUE) were calculated for each genotype at each TP of the HTP data (PH, LAI, and WL) as follows:

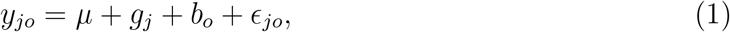

where *y*_*jo*_ is the phenotypic observation for the *j*th genotype in the *o*th block, *µ* is the intercept, *g*_*j*_ is the genotype fixed effect, *b*_*o*_ is the block random effect, and *ϵ*_*jo*_ is the model residuals. The BLUE values were used for further analysis. We calculated five spectral vegetation indices (SVI) for each genotype at each TP using the BLUE values of the WL data - NDVI (Rouse Jr et al., 1974), Green Normalized Difference Vegetation Index (GNDVI, Gitelson et al. (1996)), Triangular Greenness Index (TGI, Hunt Jr et al. (2011)), Normalized Difference Red-Edge index (NDRE, Barnes et al. (2000)), and Red-Edge inflation point (REIP, Dawson and Curran (1998)). More information on the SVI calculation is provided in Table S1.

### 2.3 Single-time point genetic analysis

We used a linear mixed model in ASREML-R version 4.2 (VSNI, UK) to estimate genomic heritability and fit genomic best linear prediction (GBLUP) for all traits on a single-time (ST) point basis. The GBLUP model was fitted as follows:

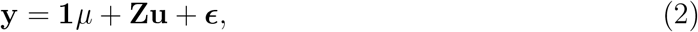

where **y** is the vector of phenotypes; **1** is the vector of ones; *µ* is the overall mean; **Z** is the incidence matrix for the random effect; 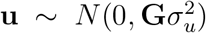 is the vector of random genotypes; **G** is the first genomic relationship matrix of VanRaden (VanRaden, 2008); and 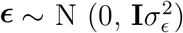 is the random residual. Genomic heritability was estimated by extracting the variance components for the additive genetic effect and residual for each trait from restricted maximum likelihood and defined as follows.

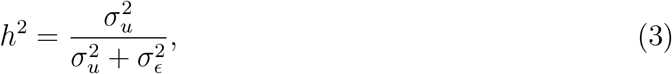

where 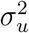 and 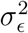 are the additive genetic and residual variances, respectively.

### 2.4 Random regression model

RR models for the longitudinal data were fit in ASREML-R as the baseline model (RR-GBLUP)

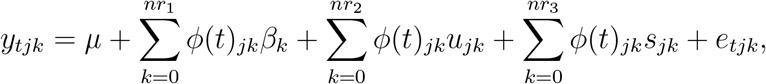

where *µ* is the grand mean; *β*_*k*_ is the *k*th order fixed regression coefficient to model the overall trend in the trait over time; *u*_*jk*_ and *s*_*jk*_ are the *k* random regression coefficients for additive genetic and permanent environmental effects of line *j*, respectively; *nr*_1_, *nr*_2_, and *nr*_3_ are the order of polynomial for the fixed and two random effects; and *e*_*tjk*_ is the random residual. The order of the Legendre polynomial for the overall mean trend (*β*) was selected by fitting only the phenotypic data for each trait. After selecting the best order of the overall fixed trend, we fit numerous polynomial functions and variance structures for the additive genetic and permanent environmental effects. For each trait, the best order for fixed and random effects was ranked based on goodness-of-prediction using Akaike’s information criterion (AIC) scores (Akaike, 1974). Genetic and permanent environmental variance components at each time point were calculated from the RR coefficients as described in Campbell et al. (2018) as 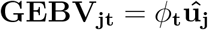, where **t**_*t*_ = *ϕ*_*tk*_ is the *t*th row vector of the matrix of Legendre polynomials at different time points (*ϕ*) for the *t*th time-point and **Ω** is the covariance matrix of RR coefficients for the genetic and permanent environmental effects. Genomic heritability for the RR model was estimated using equation (3) by adding the permanent environmental effects to the denominator part. GEBV for line *j* at time *t* were calculated as **GEBV**_**jt**_ = *ϕ*_**t**_**û**_**j**_, where *ϕ*_**t**_ is the row vector of the matrix in the order of fit of the Legendre polynomials for each trait and **û**_**j**_ is the coefficient of the additive genetic effect.

### 2.5 Genome-wide association models

Two GWAS models were used to detect genomic regions associated with longitudinal PH in sesame. The first approach was a single-marker linear mixed model regression GWAS (SM-GWAS) using the rrBLUP R package (Endelman, 2011) as follows:

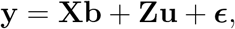

where **y** is the vector of phenotypes; **b** is the vector of fixed effects including one SNP and the top three principal components (PCs); **u** is a vector of random additive genetic effect with mean zero and variance-covariance 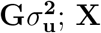 and **Z** are the known incidence matrices; and ***ϵ*** is the vector of residuals. Here, PCs and **G** were used for controlling population structure and relatedness between lines. The second approach was a GBLUP-based GWAS (GBLUP-GWAS), where we converted GEBV from either ST point or RR analysis into SNP marker effects (Wang et al., 2012) and derived their *p*-values (Aguilar et al., 2019) using BLUPF90 family software (Misztal et al., 2002). Furthermore, we performed both GWAS approaches on the RR parameter estimates (i,e., linear and quadratic slopes) to identify genomic regions associated with these variations. In both approaches, the *p*-value threshold of 2.153018*∗*10^*−*5^ was determined by calculating the number of effective independent tests (Meff) using the following formula: *p* = 1 *−* (1 *−* 0.05)^(1*/*Meff)^, where 0.05 is the desired significance level and Meff = 2,382 (Li and Ji, 2005).

### 2.6 Multi-trait linear mixed models

Multi-trait GBLUP models were fit in ASREML-R by extending the univariate model presented in equation (2) to bivariate, where **y** is the stacked vector of multi-trait phenotypes containing a secondary trait (any of the measured HTP data) and the end-of-season target trait (PH at 87 DAS and seed yield). The seed yield data collection is described in a previous study (Sabag et al., 2021), and BLUE was calculated using equation (1). The following distributions were assumed for random effects:

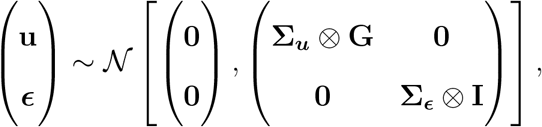

where **Σ**_***u***_ and **Σ**_***ϵ***_ refer to the genetic and residual variance-covariance matrices, respectively. Genetic correlations between the secondary and target traits were inferred from the estimated genetic parameters.

### 2.7 Cross-validation scenarios

We use several cross-validation (CV) scenarios to evaluate the prediction accuracy for the different models (Figure 1). For the RR models (Figure 1A), we used a repeated random subsampling prediction (RR-CV1) and three forecasting prediction approaches (RR-CV2, RR-CV2,1, and RR-CV2.2). RR-CV1 was used to predict the performance of new genotypes at each time point and compared to the single-time point approach (ST-CV1, Figure 1B). Forecasting approaches were used to predict the future phenotype of the same genotypes at later time points using earlier information as a training set as we increased the number of time points from 3 (RR-CV2) to 5 (RR-CV2.2).

**Figure 1:**
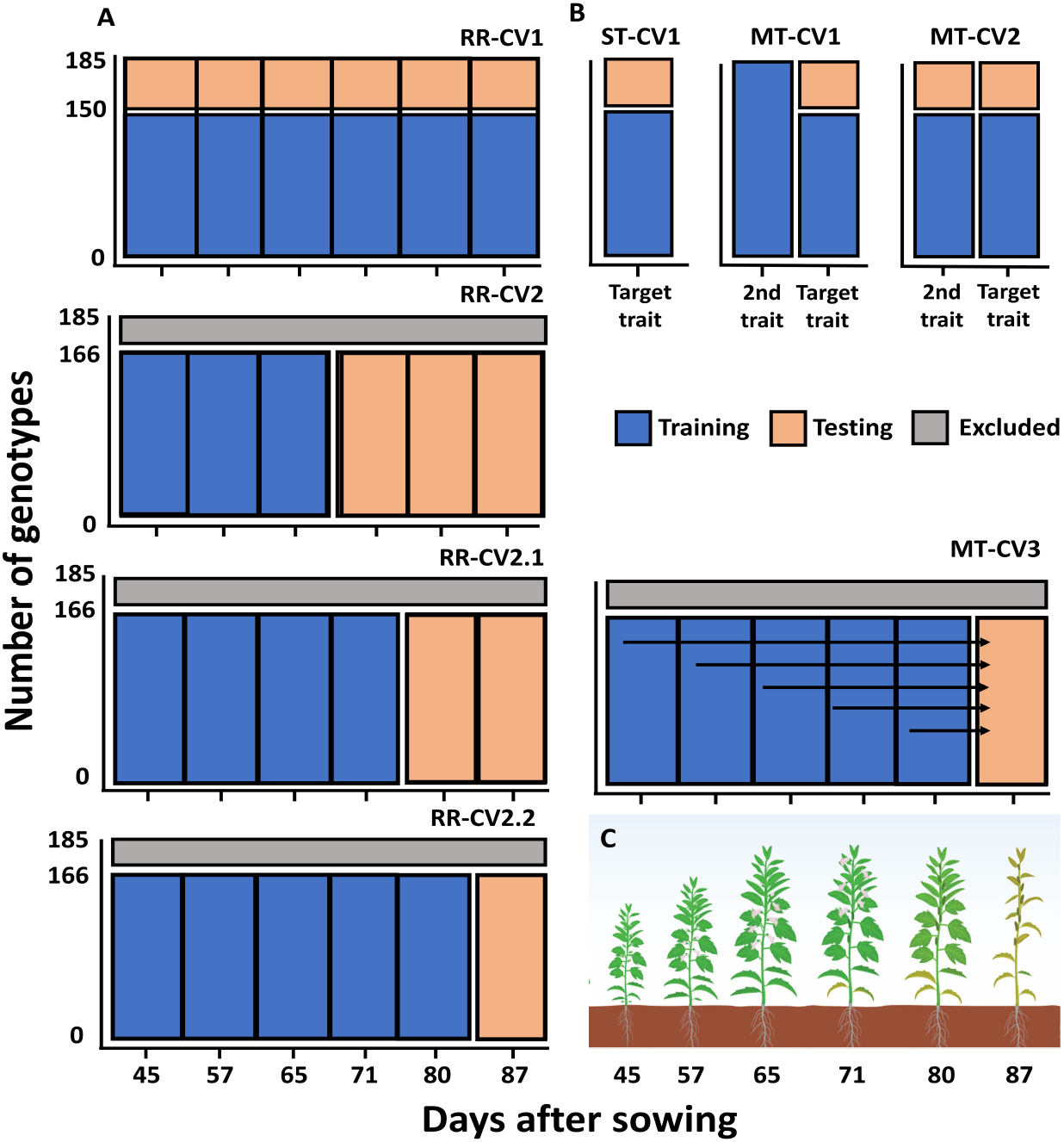
Graphical demonstration of cross-validation schemes and growth stages of sesame. Scenarios using random regression (**A**) for prediction (RR-CV1) and forecasting (RR-CV2, RR-CV2.1, and RR-CV2.2). ST-CV1 is the baseline model for a single-time point and trait, and MT-CV1, MT-CV2, and MT-CV3 are the multi-trait approaches (**B**). Growth stages of sesame range from 45 to 87 days after sowing (**C**).

The accuracy of multi-trait genomic prediction was evaluated under three scenarios (Figure 1B). In the first scenario (MT-CV1), the training set contains the secondary trait phenotype for all genotypes and only part of the target trait (seed yield). In the second scenario (MT-CV2), the same proportion was missing for the secondary trait. The last scenario (MT-CV3) is a combined approach between subsampling and forecasting prediction, where we used each of the earlier time points (45 to 80 DAS) to predict the last time point (87 DAS).

For the prediction approaches (RR-CV1, MT-CV1, and MT-CV2), the panel was randomly divided into 150 and 35 genotypes for the training and testing sets, respectively, and this process was repeated 50 times. For the forecasting models (RR-CV2, RR-CV2.1, RR-CV2.2, and MT-CV3), we randomly selected 90% of the panel for analysis (10% were excluded), and this procedure was repeated 10 times. Prediction accuracy was evaluated for all the models as the mean Pearson correlation coefficient between the observed and predicted phenotypic values of the testing set.

## 3. RESULTS

### 3.1 High-throughput phenotyping revealed diverse overall trajectories for longitudinal traits

To track sesame development during the growing season, seven longitudinal traits were measured in this study: two morphological, PH and LAI, and five SVI (Table S1). For morphological traits, the average PH of all genotypes ranged from 59 cm at 45 DAS to 163 cm at 87 DAS, while LAI values were 3.78 at 45 DAS and 4.83 at 87 DAS (Figures 2A, B, and Table S2). Modeling the whole fixed phenotypic trajectory for the morphological traits resulted in different patterns (Table S3). The best order of the Legendre polynomial fit to the fixed trajectory was 2 for PH, when it showed a steady increase from 45 to 80 DAS and then reached a plateau. On the other hand, the 4th order Legendre polynomial was the best fit to the fixed trajectory for LAI, peaking at 57 DAS and then starting to decline. For SVI, which examines various patterns along the season (Figures 2C-G, Tables S2, and S3), the values of NDVI and TGI were high at the first time point (0.89 and 6.40, respectively) and decreased toward the end of the season (0.78 and 1.54, respectively). GNDVI, NDRE, and REIP showed similar trajectories as they increased between 45 and 65 DAS and then slowly declined until 87 DAS. The best order of the Legendre polynomial for fixed trajectories for NDVI, GNDI, and TGI was 3rd order, while 4th order was the best for NDRE and REIP (Table S3).

**Figure 2:**
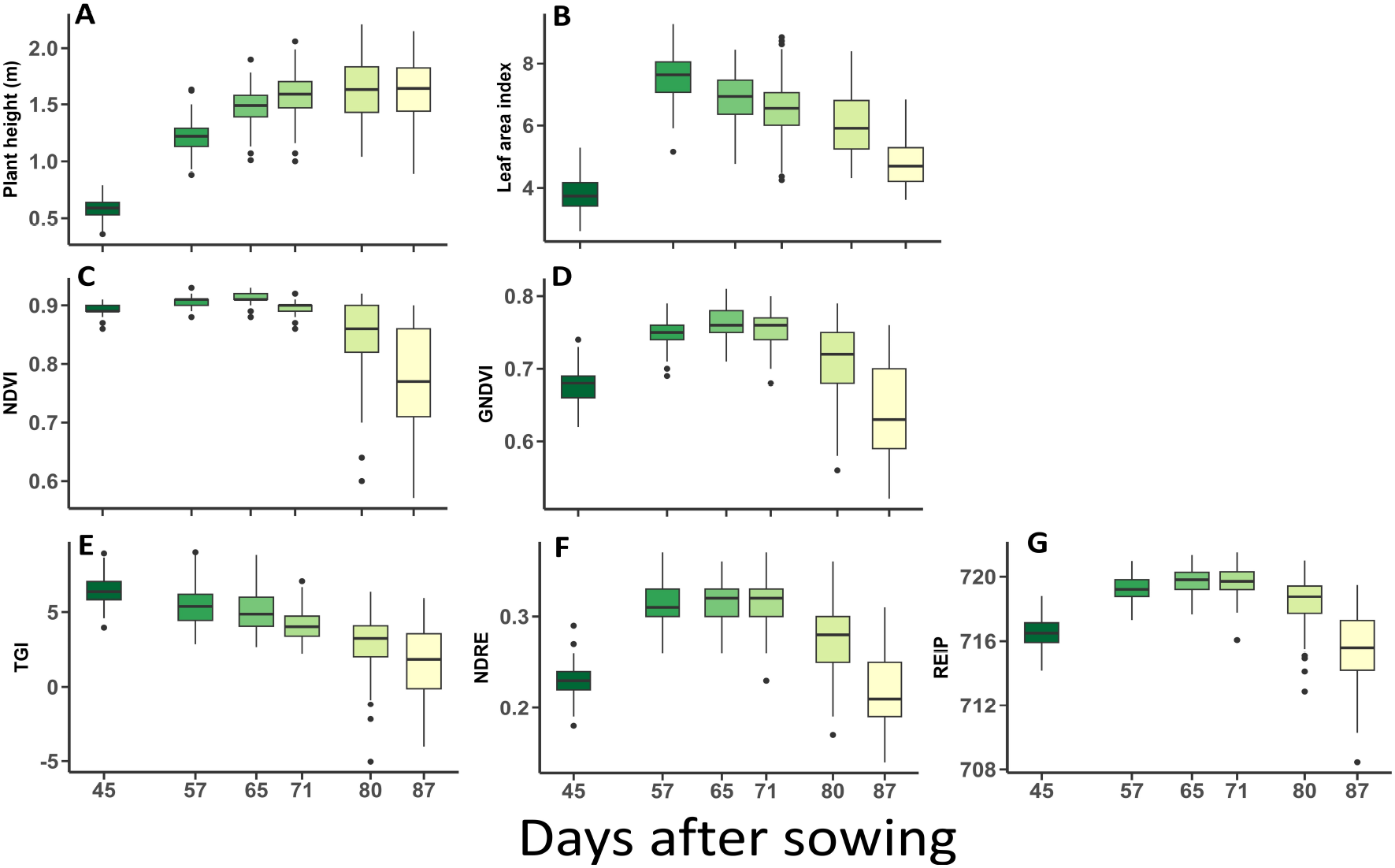
Phenotypic trajectory of longitudinal traits. Plant height (**A**), Leaf area index (**B**), Normalized Difference Vegetation Index (NDVI, **C**), Green Normalized Difference Vegetation Index (GNDVI, **D**), Triangular Greenness Index (TGI, **E**), Normalized Difference Red-Edge index (NDRE, **F**), and Red-Edge inflation point (REIP, **G**). The color gradient reflects the phenology of the sesame from greenness to senescence.

In addition, we observed changes in the coefficient of variation (CV) across time points. Overall, they tended to increase as the growing season progressed (Table S2). For example, the CV for PH was 13.92 at 45 DAS, decreased to 9.79 at 65 DAS, and increased again to 15.43 at 87 DAS.

### 3.2 Temporal genetic parameters of morphological traits and spectral indices

A primary goal of this study was to investigate the genetic components underlying the measured traits throughout the growing season using the RR framework. After determining the best model for each trait, this information was integrated into the RR model framework to identify the patterns of additive genetic and permanent environmental effects. Multiple models were fitted for each trait, including different orders of the Legendre polynomials for both genetic and environmental effects. The optimal fit was selected based on AIC scores, as shown in Table S3. For PH and LAI, the best order of fit was 2 for additive genetic effect, while the permanent environmental effects were 1 and 2, respectively. The best order of additive genetic effect for SVI was 2 for NDVI, NDRE, and REIP, while 3 for GNDVI and TGI, showing more complex patterns. The best order of fit for the permanent environmental effect of NDVI, GNDVI, and REIP was 1 and 2 for TGI and NDRE.

Variance components derived from the fitted RR model enabled the computation of genomic heritability at different time points throughout the growing season, allowing comparative analysis with the heritability estimates for ST point (Figure S2 and Table S4). The estimates of genomic heritability for PH ranged from 0.14 to 0.45 for ST and 0.25 to 0.65 for RR. Notably, the RR approach resulted in higher estimates at early and middle time points (45, 57, 65, and 71 DAS). Both models showed similar trends for LAI (Figure S2 and Table S4) when it reached its lowest values at 65 DAS (0.09 for ST and 0.23 for RR), followed by an increase to higher values. In addition, The RR approach showed higher values at earlier stages (45, 57, and 65 DAS), with the two models converging in the later stages (71, 80, and 87 DAS). For GNDVI, the ST model yielded higher estimates at 57 and 65 DAS (0.77 and 0.68, respectively) compared to 0.61 and 0.41 for RR. However, this trend reversed at 87 DAS, when RR resulted in an estimate of 0.86, higher than the 0.5 observed for ST. RR did not capture genetic variation for NDVI, REIP, and NDRE and showed consistently low genomic heritability estimates across all time points (Figure S2 and Table S4). On the other hand, the ST approach suggested the presence of genetic components for NDRE and REIP, especially at 57 DAS (0.74 and 0.87) and 65 DAS (0.75 and 0.84).

### 3.3 Time-series genetic architecture of plant height

Plant height is an important trait in sesame due to its indeterminate growth habit. Here, we used the RR framework to perform RR-GBLUP and GWAS and compared it with the ST point approach.

#### 3.3.1 Prediction and forecasting

Predicting new genotypes at each time point (Figure 1A, RR-CV1) resulted in accuracies between 0.27 and 0.65 for RR-GBLUP and 0.33 to 0.65 for ST-GBLUP (Figure 3A and Table S5). ST-GBLUP performed better than RR-GBLUP at 45 (0.33 vs. 0.27), 57 (0.32 vs. 0.38), and 65 (0.46 vs. 0.49) DAS, while the models showed similar performance at later time points (Table S5). For the forecasting scenarios, RR-CV2, RR-CV2.1, RR-CV2.2, and MT-CV3 enabled the prediction of future phenotypes using the earlier time points as the training set (Figure 1A). Overall, prediction accuracy increased as we used closer time points for the training sets. In RR-CV2, when we used three earlier time points as the training set, the accuracy was 0.78, 0.46, and 0.17 for 71, 80, and 87 DAS, respectively, and increased to 0.84 (80 DAS) and 0.73 (87 DAS) in the RR-CV2.1 scenario (Figure 3B and Table S5). The highest accuracy was achieved with RR-CV2.2 for 87 DAS (0.86). In the last scenario (MT-CV3), we investigated which time point was best at predicting the last time point (87 DAS) in a multi-trait model (Figure 1A). This analysis performed worst when we used the 45 DAS phenotype, and the prediction accuracy was 0.17. However, we observed a steady improvement thereafter from 57 (0.62) to 80 DAS (0.90) (Figure 3C and Table S5). Accuracy using the 80 DAS phenotype outperformed all other models for predicting end-of-season PH, followed by 71 (0.82) and 65 DAS (0.76).

**Figure 3:**
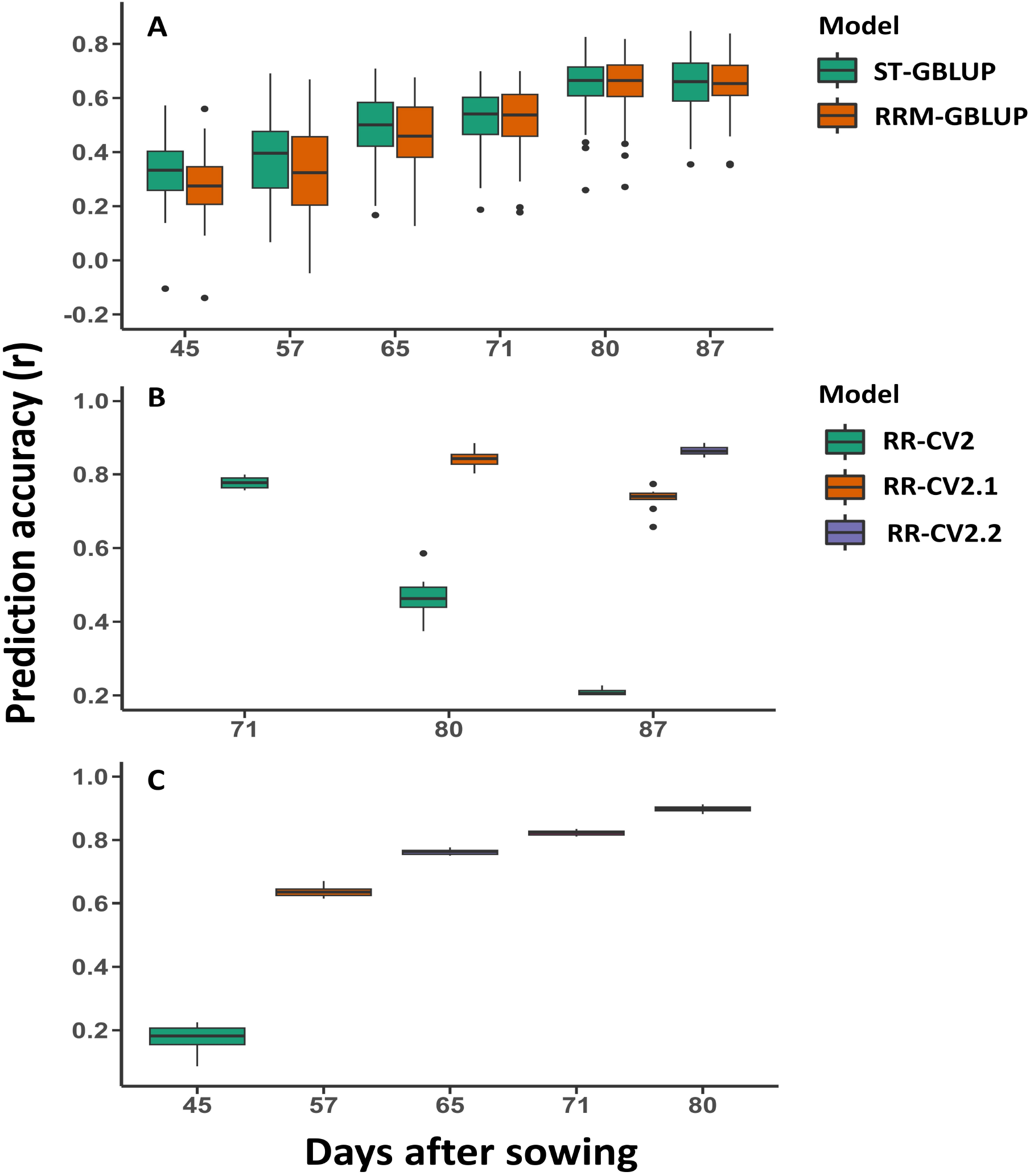
Plant height prediction accuracy for different models. Comparison between singletime point and random regression genomic best linear unbiased prediction to predict new genotypes at each time point (**A**). Forecasting future phenotypes (**B**) using three (RR-CV2), four (RR-CV2.1), and five (RR-CV2.2) earlier time points as training sets, respectively. Multi-trait models (MT-CV3, **C**) using earlier time points (45 to 80 days after sowing) to forecast end-of-season plant height phenotype (87 days after sowing). Prediction accuracy was obtained by Pearson correlation between genomic estimated breeding values and observed values in the testing sets.

#### 3.3.2 Longitudinal GWAS

Investigating the genetic architecture of a trait over time helps to identify QTL that contributes to its variation over the growing season. In this study, two GWAS approaches, SM-GWAS and GBLUP-GWAS, were used to identify QTL for PH over the growing season. Notably, GBLUP-GWAS, which uses GEBVs from ST and RR-GBLUP analyses, identified more significant QTLs compared to SM-GWAS.

##### Single-time point approach

With SM-GWAS, three QTLs on linkage groups (LGs) 6, 8, and 11 reached the threshold at 80 and 87 DAS (Figure 4A). The QTL on LG11 was significant at both dates (*p*-values = 5.27 and 4.98, respectively), while the QTLs on LG8 at 80 DAS and LG6 at 87 DAS were slightly below the threshold (*p*-values = 4.42 and 4.61, respectively). Since the *p*-values from GBLUP-GWAS were relatively low (highly significant), we selected the 10 most significant SNPs at each time point as potential QTLs for PH. For example, using the ST point GBLUP-GWAS approach, we discovered the top 10 significant SNPs at 57 DAS located in six different LGs (Figure 4B). The SNP with the lowest *p*-value was mapped to LG6, and four SNPs were located on LG9, which were also significant at 65 DAS. On LG12, we found six significant SNPs at 65 and 71 DAS, spanning the genomic interval of *∼*235,000 base pairs. The SNP on LG11, which was also found to be significant in SM-GWAS, was among the top 10 SNPs from 65 DAS to the end of the season (87 DAS), along with the LG8 SNP, which was found to be significant at the three later time points. In addition, SNPs located on LG6 and LG10 were of high magnitude at 80 and 87 DAS, suggesting their potential role as QTLs for the end-of-season phenotype.

**Figure 4:**
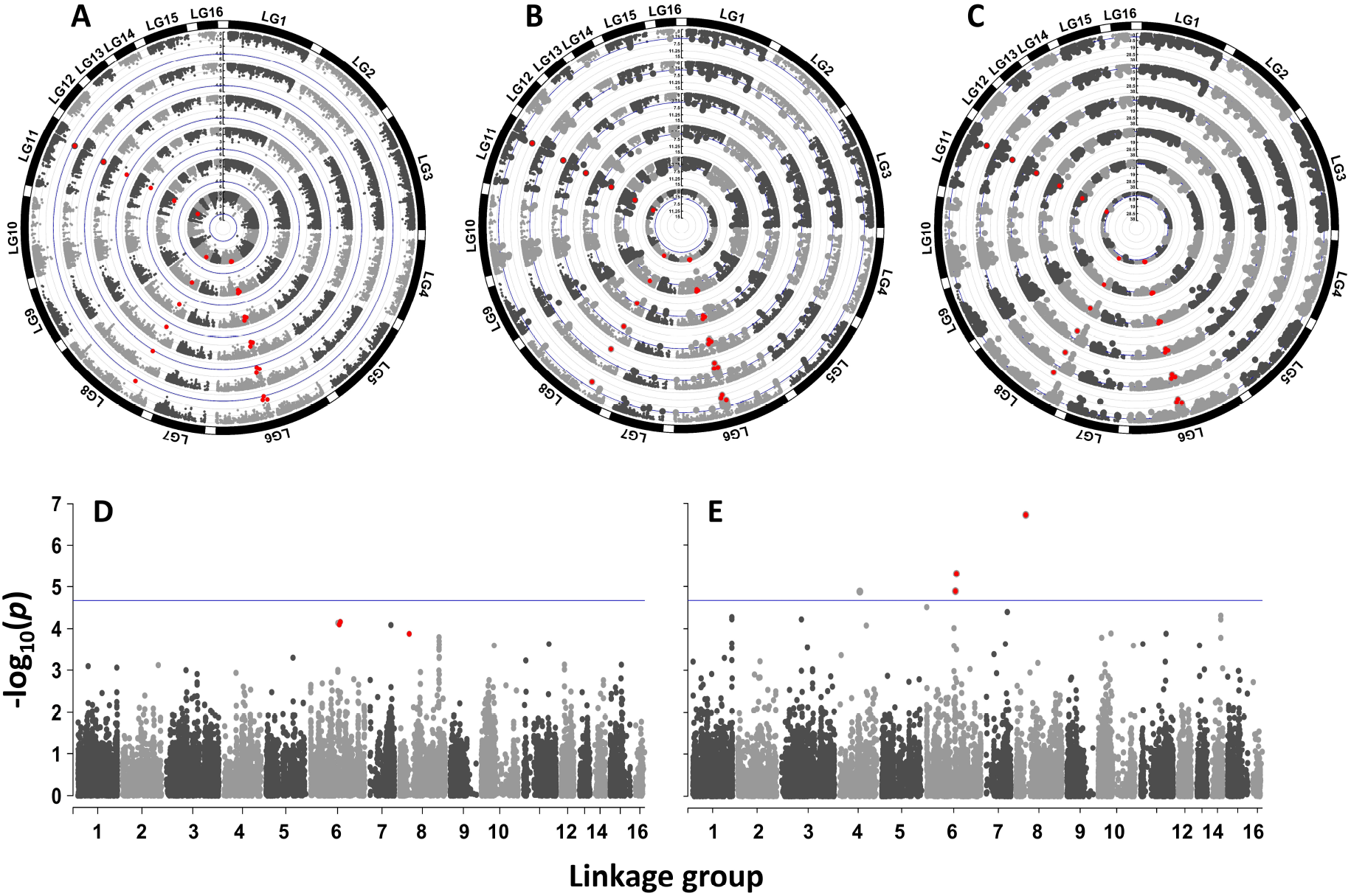
Genome-wide association study (GWAS) results for plant height and random regression genetic parameters. Circular Manhattan plots for the six time points, from the inner circle (45 days after sowing) to the outer circle (87 days after sowing). Single-marker linear mixed model regression GWAS (**A**), genomic best linear unbiased prediction-based GWAS for single-time and random regression models (**B** and **C**). Manhattan plots for the linear slope using the two GWAS models (**D** and **E**). Red highlighted points are genetic markers that were found to be significant for both traits in both approaches.

##### Time-series approach

The RR-based GBLUP-GWAS allowed us to consider the whole phenotype trajectory by incorporating the GEBV from the RR into the GBLUP-GWAS model (Figure 4C). Three SNPs located at LG3, 7, and 8 were uniquely associated with the trait at 45 DAS, and two SNPs at LG4 and 5 were found to be significant at 45 and 57 DAS. These SNPs were only discovered in the RR GBLUP-GWAS model at the earlier time points. Additionally, two novel SNPs on LG2 and LG12 had a large effect at the three earlier time points, while the QTL on LG9 (also mapped in other models) was significant up to 71 DAS. Towards the end of the season, at 80 and 87 DAS, the three QTLs identified in the other two models on LGs 6, 8, and 11 were significant in the RR approach, along with newly identified SNPs on LGs 1, 6, 7, 8, and 10 (Figure 4C). In addition, we leveraged the phenotypic trajectory to perform GWAS on RR parameters, including linear and quadratic slopes. Using the two GWAS approaches, we mapped significant QTLs on LG6 and LG8 for the linear slope (Figure 4D and E) consistent with their significance for PH in all other models. In the final step, we examined the differences between QTLs affecting the trait variation at an ST point and QTLs affecting the linear slope (i.e., growth rate) by focusing on the QTLs at LG6 and LG11, respectively. The observed variation for the linear slope ranged from -0.15 to 0.18 (Figure 5A), and the LG6 QTL distinguished between the high (C allele) and low (T allele) linear slope (Figure 5B). Furthermore, Figure 5C-D shows the dynamics of PH in the diversity panel along the growing season, highlighting the difference in linear growth rate between genotypes harboring high and low alleles for the LG6 QTL (Figure 5C) and the mean differences between the LG11 QTL across the growing season (Figure 5D). These results illustrated the ability of RR to detect QTLs influencing both ST points and growth rate variation.

**Figure 5:**
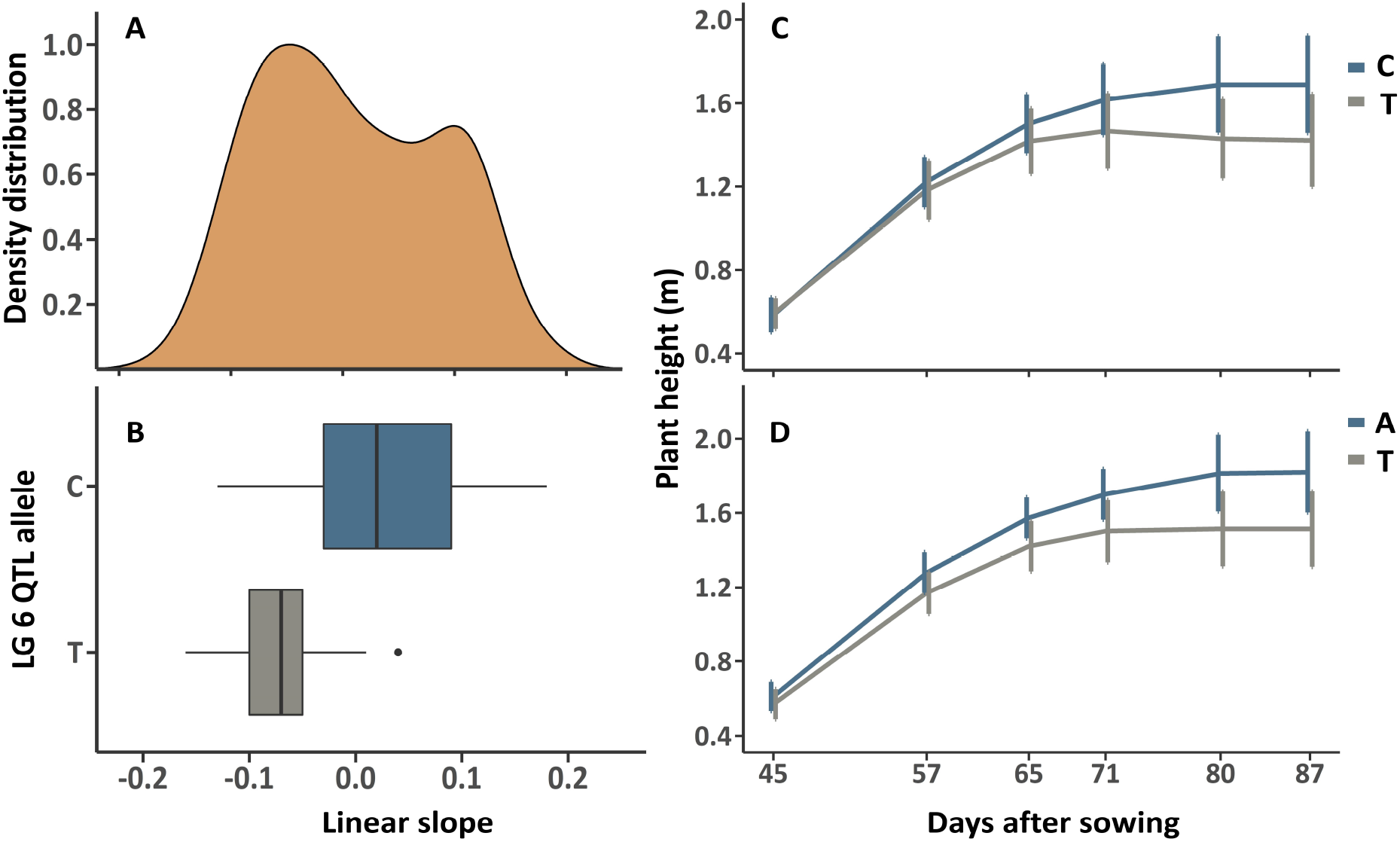
Genome-wide association analysis of the linear slope obtained from the random regression model. Density distribution for linear slope values in the sesame diversity panel (**A**) and allelic configuration of linkage group (LG) 6 quantitative trait loci affecting the linear slope (**B**). Plant height development of genotypes harboring different alleles for LG6 and LG11 quantitative trait loci affecting both linear slope and mean variation (**C**) and mean variation only (**D**), respectively.

### 3.4 Multi-trait prediction on seed-yield

HTP provides a framework to record traits over time. It allows HTP-derived traits to be used as secondary traits to predict a target trait, typically an end-of-season phenotype. Here, we used both temporal morphological traits and SVI to predict seed yield. First, we obtained the phenotypic and genetic correlation for each trait at each time point with seed yield. For the morphological traits (PH and LAI), the phenotypic and genetic correlations showed a similar pattern, turning from positive in the earlier stages to negative in the later stages (Figure S3 and Table S5). The phenotypic and genetic values ranged from 0.28 and 0.11 to -0.3 and -0.73 for PH, 0.02 and 0.19 to -0.27 and -0.49 for LAI, respectively. NDVI and TGI showed mostly negative values for both types of correlations, as the lowest value for the phenotypic correlation was -0.22 and -0.38 at 65 and 71 DAS, respectively. Both traits showed a similar negative genetic correlation (-0.75 for NDVI and -0.77 for TGI) at 71 DAS. Notably, the pattern of correlations for GNDVI, NDRE, and REIP showed positive values for both correlation types at the first four time points and then showed a steep decline to negative values at 80 and 87 DAS (Figure S3 and Table S5). Furthermore, we leveraged the correlations between temporal secondary traits and seed yield to perform multi-trait GBLUP and investigate which trait at which time point can enhance seed yield prediction. Two cross-validation schemes (MT-CV1 and MT-CV2, Figure1B) were used to evaluate the prediction accuracy of these models and compare them with the single-trait GBLUP of seed yield (ST-CV1). Overall, we observed a slight improvement when using the phenotype from the earlier time points rather than the later ones (Figure 6 and Table S6). At 45 and 57 DAS, NDRE and REIP showed better prediction accuracy (0.43 and 0.44, respectively, for both traits) than the single-trait GBLUP (0.42) using the MT-CV1 scenario (Figure 6A and B), while observed no improvement for the MT-CV2 scenario. PH showed some improvement at 45 DAS (0.44) and then declined to 0.35 and 0.34 at 57 and 65 DAS, respectively, with the MT-CV1 scenario. Interestingly, in the MT-CV2 scenario, the prediction accuracy for PH increased to 0.43 at 57 and 65 DAS (Figure 6B and C). No improvement in LAI was observed at the first three time points in either scenario. We found no improvement for GNDVI and TGI at 45 DAS, but at later time points (57 and 65 DAS), the accuracy increased to 0.44 (for GNDVI) and 0.43 (for TGI), respectively, in the MT-CV1 scenario. For the later time points (71, 80, and 87 DAS), we observed no evidence of improvement using the secondary traits, except for PH and REIP (0.43) at 71 DAS in the MT-CV2 scenario.

**Figure 6:**
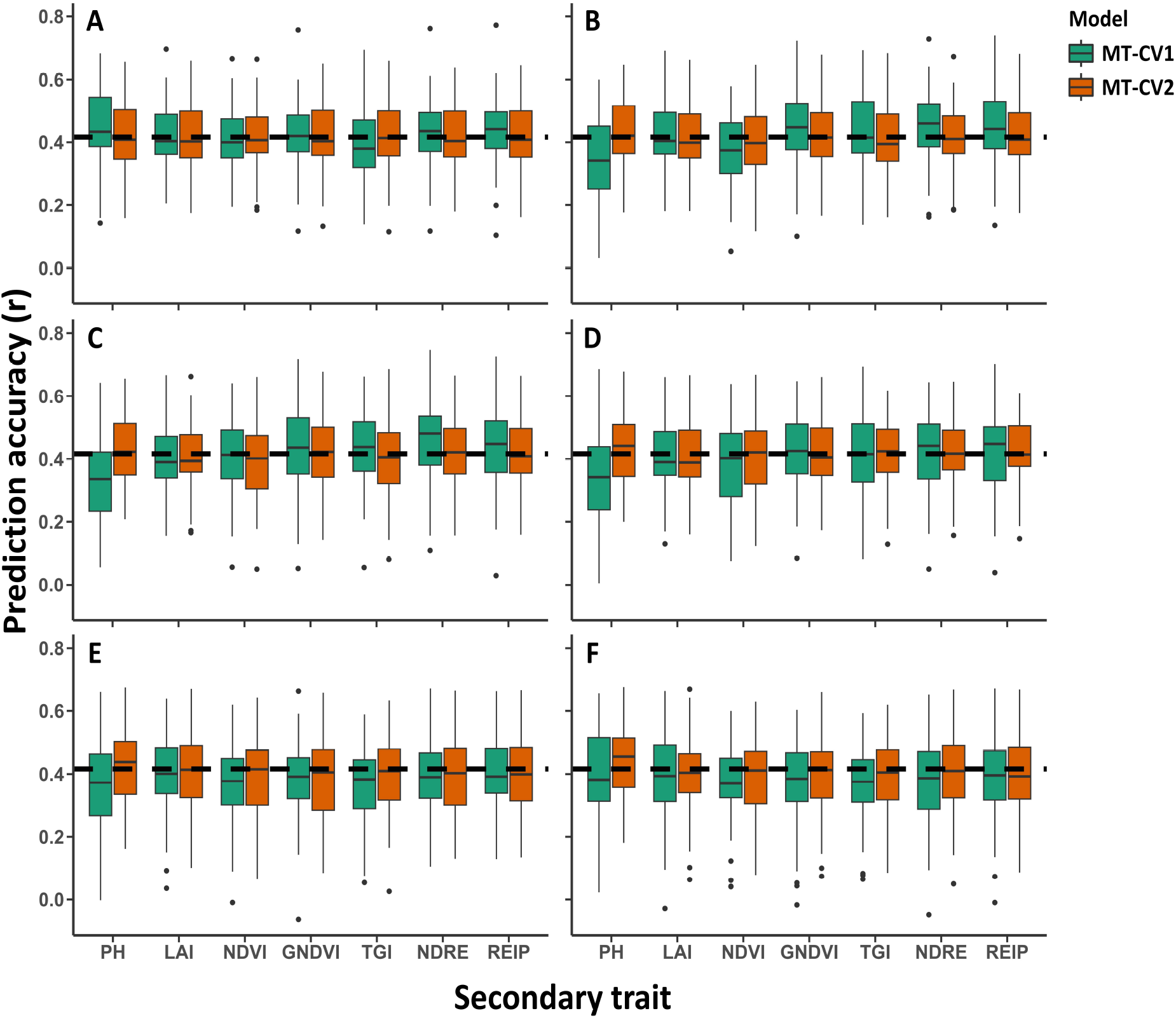
Multi-trait genomic best linear unbiased prediction accuracy incorporating temporal phenotypes and seed yield at each time-point using the two cross-validation scenarios at 45 (**A**), 57 (**B**), 65 (**C**), 71 (**D**), 80 (**E**), and 87 (**F**) days after sowing. The dashed black line represents the mean prediction accuracy for a single-trait genomic best linear unbiased prediction analysis for seed-yield.

## 4. DISCUSSION

Ensuring food security relies on integrating new crops with climate change resilience and high nutritional values (Ribaut and Ragot, 2019). Orphan or neglected crops, such as sesame, offer useful genetic resources for identifying new alleles and traits contributing to climate adaptability and nutritional properties. Thus, major efforts should be made to develop efficient breeding programs to rapidly increase the genetic gain in the crop of interest. Since sequencing and genomic technologies are not limiting factors, phenotyping of large plant populations is still a bottleneck in plant breeding and genetics studies (Araus et al., 2018; Song et al., 2021).

### 4.1 Time-series analysis promotes the detection of sesame development along the growing season

An HTP approach was developed for the sesame diversity panel to explore its potential for investigating the genetic architecture of several longitudinal functional traits during the growing season and their contribution to final seed yield. Focusing on the reproductive phase, we were able to track the development of two morphological traits and five SVI that project canopy status (Table S1). These phenotypic trajectories captured by the HTP platform were consistent with our knowledge of sesame development. For example, PH consistently increased until the end of the season, and LAI was high at the beginning of the reproductive stage and then decreased toward the end. SVI can assist by tracking the plant’s physiological status (Sarić et al., 2022). The use of NDVI and NDRE has been studied in other sesame populations (Dong et al., 2021; Petsoulas et al., 2022), and they reported similar results when these two indices reached the highest values at the peak of the reproductive stage (57 to 70 DAS) and then decreased as the plants started senescence. This suggests that our HTP platform is reliable for monitoring longitudinal traits in sesame, although additional analysis is required to dissect these traits in the vegetative phase.

### 4.2 Genetic control of longitudinal trait in sesame

Modeling the additive genetic effect through RR models revealed that most traits have a less complex growth pattern in the genetic part (Table S3). For example, the best fit for most traits was second order, except for TGI and GNDVI, which had a third-order best fit. Estimation of genomic heritability showed the temporal genetic contribution of phenotypic variation for each trait calculated on an ST point basis and from the RR. Comparing these two approaches (Figure S2 and Table S4), we observed that for some of the traits, RR produced higher genomic heritability estimates for PH and LAI, especially in the earlier time points, while for NDRE and NDVI, the ST point approach showed higher values than RR approach. For GNDVI, TGI, and NDVI, the results were almost the same for the ST and RR approaches. Moreover, genomic heritability estimates varied between time points, suggesting that the environmental effect also varied between time points, which was also observed in soybean (Freitas Moreira et al., 2021). In addition, measuring traits at denser time points may be beneficial for understanding heritable genetic components and result in a smoother trend line, as examined in rice (Campbell et al., 2018).

### 4.3 Random regression analysis highlights the temporal genetic architecture of plant height

Plant height is a key trait in indeterminate crops, such as sesame, affecting the whole plant architecture, seed yield, and yield components (Kim et al., 2022; Kuzbakova et al., 2022). Several studies have been conducted in sesame to assess the genetic architecture and optimal prototype of PH to maximize seed yield (Wei et al., 2015; Wang et al., 2016; Miao et al., 2020). Here, we used time-series analysis of PH throughout the growing season and employed GWAS and genomic prediction using the RR and ST point approaches. Comparing the prediction accuracy of these two approaches in the RR-CV1 scenario (Figure1A) showed nearly identical values (Figure 3A), which is in contrast to our hypothesis that RR can capture more additive genetic variance by accounting the entire phenotypic trajectory. In addition, the prediction accuracy of the two approaches reached higher values at the last time point (80 and 87 DAS), suggesting that PH is more influenced by genetics in these last stages of the growing season. The forecasting approach allowed us to predict the phenotype of the same genotypes in later periods using the earlier ones as a training set (Figure 3B). Overall, we found that prediction accuracy increased when we used closer time points as training sets (from 3 to 5), and similar results were found in rice under constrained water regimes (Momen et al., 2019). Moreover, coupling the 87 DAS phenotype with any other earlier DAS phenotypes in a multi-trait model (scenario MT-CV3) showed a positive result (Figure 3C), implying that around 71 DAS we can achieve a reliable prediction of PH at the end of the season. Since sesame is an indeterminate crop plant, forecasting the future phenotype can assist breeders in decision-making and selecting the desired phenotype at earlier stages.

Longitudinal GWAS was performed using two approaches: SM-GWAS and GBLUP-GWAS. The continuous mapping discovered numerous QTLs controlling PH along the growing season (Figure 4). This approach presented major advantages over the traditional one of mapping the finite phenotype at the end of the season. Using the same dataset was used, no significant genomic signal was detected (Sabag et al., 2021). Overall, GBLUP-GWAS (in both ST points and RR) was able to detect more significant regions (Figure 4A-C), suggesting that PH is controlled by minor QTLs along the growing season. In GBLUP-GWAS, we were able to map two notable QTLs on LGs 9 and 12 at early-mid DAS, which were located within the QTL interval detected in the sesame F_2_ population for PH and internode length (Teboul et al., 2020). In addition to detecting common QTLs, RR-based GBLUP-GWAS tended to result in larger sizes, as reflected in *p*-values. For example, the highest *−log*_10_(*p −* value) of this model was 36.56 compared to 14.23 of ST-based GBLUP-GWAS, consistent with Campbell et al. (2019), which found a larger SNP effect using RR in rice. The RR parameters, especially the linear slope, reflect the growth rate of each genotype. We obtained variation in linear slope values in our diversity panel (Figure 5A), and we were also able to detect QTLs associated with the linear slope on LGs 4, 6, and 8 (Figure 4E). In our panel, the QTLs on LGs 4 and 8 distinguished between genotypes harboring the major allele and the heterozygous genotype, so we could not assess their magnitude in the downstream analysis. The LG6 QTL, also found to be associated with PH across time, differs between genotypes with high and low linear slope values (Figure 5B and D) and appears to serve as an important genetic component regulating PH development in sesame. There are two candidate genes (LOC105164490 and LOC105164477) near the significant SNPs that encode cellulose synthase-like protein G3 and xyloglucan endotransglucosylase/hydrolase protein 30, respectively, which involved in stem cell wall composition, plant height, and cell elongation in other crops (Jiménez et al., 2006; Ding et al., 2015; Hyles et al., 2017).

### 4.4 Leveraging secondary traits to enhance seed-yield prediction

Seed yield is an interrelated and complex result of multivariate relationships and compensations between various yield components and crop characteristics, as well as their interactions with management and environmental cues during the growing season. HTP platforms provide noninvasive approaches to measure temporal traits and monitor crop development at various stages. Here, we leveraged the correlation between the seven temporal traits (PH, LAI, and five SVI) and seed-yield to perform multi-trait genomic prediction (MT-GBLUP). Overall, there was no statistically significant improvement in either scenario (MT-CV1 and MT-CV2, 1B) compared to the single-trait analysis for seed yield. However, it is worth noting that when using NDRE and REIP as secondary traits, improvements were achieved from 45 to 71 DAS, and also for GNDVI and TGI at 57 and 65 DAS in the MT-CV1 scenario. These results are consistent with the high genomic heritability estimates and correlations with seed yield at these stages (Figures S2 and S3). Similar results were obtained in wheat and maize (Sun et al., 2017; Anche et al., 2020) when using SVI to improve seed yield, especially in the MT-CV1 scenario. The lack of statistically significant improvement in the multi-tait model could be because both morphological traits and SVI had moderate to high phenotypic correlations with flowering date, especially in the later time points (data not shown). Previous studies on this data set (Sabag et al., 2021) found a negative correlation between flowering date and yield (r = -0.54) and a high heritability for flowering date (*H*^2^ = 0.97). Considering these findings, it may be beneficial to perform a two-stage model, first estimating flowering date using remote sensing approaches and then predicting seed yield as demonstrated for soybean (Toda et al., 2021).

## 5. CONCLUSIONS

The HTP platform developed in this study demonstrates its reliability in estimating sesame development throughout the growing season. When coupled with RR, it proves valuable in modeling and unraveling the genetic contributions of temporal-continual traits. Notably, RR can be considered as a useful tool for identifying new QTLs associated with PH, with higher magnitudes compared to alternative approaches. This versatile approach can be extended to explore the genetic architecture in time-series data for other traits. Furthermore, the integration of these temporal traits into genomic prediction models provides some improvement in seed yield prediction, highlighting the potential of HTP in this context. While our findings provide valuable insights, further studies are essential to unravel the inheritance components of longitudinal traits in different environments. Additionally, investigating genotype-by-environment interactions, including both morphological and spectral traits, is crucial to fully realize the potential of HTP in sesame genetic research and breeding.

## Abbreviations

BLUE: Best linear unbiased estimates
CV: Cross-validation
DAS: Days after sowing
GBLUP: Genomic best linear unbiased prediction
GEBV: Genomic estimated breeding value
GNDVI: Green Normalized Difference Vegetation Index
GWAS: Genome-wide association study
HTP: High-throughput phenotyping
LAI: Leaf area index
MT: Multi-trait
NDRE: Normalized Difference Red-Edge index
NDVI: Normalized Difference Vegetation Index
PH: Plant height
QTL: Quantitative trait loci
REIP: Red-Edge inflation point
RR: Random regression
ST: single-time
SVI: Spectral vegetation indices
TGI: Triangular Greenness Index
TP: Time point
WL: Wavelengths

## AUTHOR CONTRIBUTIONS

**Idan Sabag:** Conceptualization; data curation; formal analysis; investigation; methodology; visualization; writing—original draft; writing—review and editing. **Ye Bi**: Formal analysis; writing—review and editing. **Maitreya Mohan Sahoo:** Methodology; analysis; writing—review and editing. **Ittai Herrmann:** Conceptualization; methodology; funding acquisition; writing—review and editing. **Gota Morota:** Conceptualization; formal analysis; funding acquisition; project administration; supervision; writing—review and editing. **Zvi Peleg:** Conceptualization; data curation; formal analysis; funding acquisition; investigation; project administration; supervision; visualization; writing—review and editing.

## ACKNOWLEDGMENTS

This research was supported by a Research Grant from BARD, the United States - Israel Binational Agricultural Research and Development Fund (No. IS-5400-21). I.S. is indebted to the Samuel and Lottie Rudin Scholarship Foundation.

## CONFLICT OF INTEREST STATEMENT

The authors declare no conflicts of interest.

## Notes

### Competing Interest Statement

The authors have declared no competing interest.

