## Supplementary material for "Leveraging genomics and temporal high-throughput phenotyping to enhance association mapping and yield prediction in sesame": SI data

### Supplementary Materials

### Methods

#### Leaf area index estimation

We estimated the leaf area index (LAI) for individual plots from their canopy hyperspectral reflectance. Using a dataset of 120 samples with their averaged canopy reflectance acquired along two independent time points (45 and 57 Days after sowing) and measured LAI values, we developed regression models by training their spectral features using the genetic algorithm-based partial least squares regression (GA-PLS regression). The model was run for 100 iterations, consisting of 76 randomly chosen samples for calibration and 20 samples for cross-validation. Each iteration selected non-repetitive samples for the process of training that minimized model overfitting due to sampling biasedness. The trained model was developed by averaging the normalized PLS coefficients across all wavelengths for 100 iterations. The model was subsequently used to independently validate 24 unseen samples of the dataset. It was observed that the GA-PLS model had robustness and consistently performed for training and cross-validation validation datasets with the best coefficient of regression (R2) of 0.9 and 0.81 and root mean squared errors (RMSEs) of 0.71 and 1.08, respectively. The normalized coefficients and the feature selection histograms emphasized the contribution of 720, 930, and 980 nm for the spectral estimation of LAI. Continuous features identified at 700-750, 900, and 970-1000 nm can be interpreted as a combined effect of leaf chlorophyll and water content diagnostic to the LAI estimation (Curran, 1989; Gamon et al., 1995; Knyazikhin et al., 1998; Sellers, 1985; Sims and Gamon, 2003). The developed PLS model was further used to estimate the LAI values for the remaining 5550 plot samples to study the time-series trends from 45 to 87 days after sowing.

### Tables

Table S1: Summary of vegetation indices calculated from the hyperspectral data. Nanometer and reflectance are represented by nm and R, respectively.

| Index | Formula |
|---|---|
| Normalized Difference Vegetation Index (NDVI) | $\frac{801_R - 670_R}{801_R + 670_R}$ |
| Green Normalized Difference Vegetation Index (GNDVI) | $\frac{801_R - 550_R}{801_R + 550_R}$ |
| Triangular Greenness Index (TGI) | -0.5 $\times((670_{nm} - 480_{nm}) \times (670_R - 550_R))$ - $((670_{nm} - 550_{nm}) \times (670_R - 480_R))$ |
| Normalized Difference Red-Edge Index (NDRE) | $\frac{790_R - 719_R}{790_R + 719_R}$ |
| Red-Edge Inflection Point (REIP) | 700+40 $\frac{((670_R + 781_R)/2) - 700_R}{(740_R - 700_R)}$ |

Table S2: Mean, standard error (SE), and coefficient of variation (CV) of the longitudinal traits at each time point.

| Days after sowing | Trait | Mean | SE | CV(%) |
|---|---|---|---|---|
| 45 | Plant height | 59 | 8.18 | 13.92 |
| | Leaf area index | 3.78 | 0.56 | 14.39 |
| | NDVI | 0.89 | 0.01 | 0.93 |
| | GNDVI | 0.68 | 0.02 | 3.45 |
| | TGI | 6.40 | 0.81 | 12.69 |
| | NDRE | 0.23 | 0.02 | 8.43 |
| | REIP | 695.72 | 0.19 | 0.03 |
| 57 | Plant height | 121 | 12.10 | 10.03 |
| | Leaf area index | 7.56 | 0.70 | 9.25 |
| | NDVI | 0.91 | 0.01 | 0.89 |
| | GNDVI | 0.75 | 0.02 | 2.55 |
| | TGI | 5.38 | 1.16 | 21.57 |
| | NDRE | 0.31 | 0.02 | 6.87 |
| | REIP | 719.27 | 0.74 | 0.10 |
| 65 | Plant height | 148 | 14.47 | 9.79 |
| | Leaf area index | 6.89 | 0.76 | 10.98 |
| | NDVI | 0.91 | 0.01 | 1.11 |
| | GNDVI | 0.76 | 0.02 | 2.36 |
| | TGI | 4.99 | 1.23 | 24.66 |
| | NDRE | 0.32 | 0.02 | 6.71 |
| | REIP | 719.8 | 0.75 | 0.10 |
| 71 | Plant height | 158 | 18.10 | 11.47 |
| | Leaf area index | 6.56 | 0.85 | 13.02 |
| | NDVI | 0.90 | 0.01 | 1.37 |
| | GNDVI | 0.76 | 0.02 | 2.49 |
| | TGI | 4.13 | 0.92 | 22.29 |
| | NDRE | 0.32 | 0.02 | 6.91 |
| | REIP | 719.69 | 0.77 | 0.11 |
| 80 | Plant height | 163 | 24.47 | 15.05 |
| | Leaf area index | 6.03 | 1.00 | 16.53 |
| | NDVI | 0.85 | 0.06 | 6.65 |
| | GNDVI | 0.71 | 0.05 | 6.35 |
| | TGI | 2.89 | 1.71 | 59.2 |
| | NDRE | 0.27 | 0.04 | 13.10 |
| | REIP | 718.45 | 1.35 | 0.19 |
| 87 | Plant height | 163 | 25.00 | 15.43 |
| | Leaf area index | 4.83 | 0.80 | 16.47 |
| | NDVI | 0.78 | 0.09 | 10.99 |
| | GNDVI | 0.64 | 0.06 | 9.32 |
| | TGI | 1.54 | 2.31 | 150.20 |
| | NDRE | 0.22 | 0.04 | 18.34 |
| | REIP | 715.50 | 2.17 | 0.30 |

Table S3: Model selection for random regression models for the longitudinal traits. For each trait, the numbers represent the order of fit of the Legendre polynomials for fixed (F), additive genetic (G), and permanent environmental (PE) effects. $AIC_F$ and $AIC_{RR}$ represent Akaike's information criterion ranking values for fixed and random regression models, respectively.

| Trait | F | G | PE | $AIC_F$ | $AIC_{RR}$ |
|---|---|---|---|---|---|
| Plant height | 2 | 2 | 1 | -624.85 | -3711.681 |
| Leaf area index | 4 | 2 | 2 | 2633.035 | 57.15 |
| NDVI | 3 | 2 | 1 | -3859.97 | -8027.649 |
| GNDVI | 3 | 3 | 1 | -4316.72 | -7582 |
| TGI | 3 | 3 | 2 | 3984.83 | 856.42 |
| NDRE | 4 | 2 | 2 | -4783.62 | -7233.175 |
| REIP | 4 | 2 | 1 | 3604.476 | 687.08 |

Table S4: Genetic parameters of the longitudinal traits for each time point. $h^2_{TP}$ and $h^2_{RR}$ denote genomic heritability estimates in single time point basis and random regression models, respectively. $r$ and $r_g$ are the phenotypic and genetic correlations between the longitudinal traits at each time point with seed yield, respectively.

| Days after sowing | Trait | $h^2_{TP}$ | $h^2_{RR}$ | $r$ | $r_g$ |
|---|---|---|---|---|---|
| 45 | Plant height | 0.31 | 0.40 | 0.28 | 0.11 |
| | Leaf area index | 0.24 | 0.29 | 0.02 | 0.19 |
| | NDVI | 0.19 | 0.12 | 0.05 | -0.08 |
| | GNDVI | 0.42 | 0.61 | 0.10 | 0.34 |
| | TGI | 0.59 | 0.76 | -0.06 | -0.24 |
| | NDRE | 0.34 | 0.25 | 0.15 | 0.47 |
| | REIP | 0.37 | 0.06 | 0.15 | 0.45 |
| 57 | Plant height | 0.14 | 0.41 | 0.12 | -0.46 |
| | Leaf area index | 0.17 | 0.32 | 0.06 | 0.11 |
| | NDVI | 0.15 | 0.01 | -0.18 | -0.56 |
| | GNDVI | 0.77 | 0.41 | 0.29 | 0.37 |
| | TGI | 0.84 | 0.40 | -0.32 | -0.44 |
| | NDRE | 0.74 | 0.19 | 0.28 | 0.37 |
| | REIP | 0.87 | 0.02 | 0.31 | 0.37 |
| 65 | Plant height | 0.20 | 0.65 | 0.02 | -0.57 |
| | Leaf area index | 0.09 | 0.23 | 0.04 | 0 |
| | NDVI | 0.22 | 0.07 | -0.22 | -0.48 |
| | GNDVI | 0.68 | 0.41 | 0.33 | 0.59 |
| | TGI | 0.76 | 0.73 | -0.38 | -0.61 |
| | NDRE | 0.75 | 0.12 | 0.36 | 0.61 |
| | REIP | 0.84 | 0.03 | 0.37 | 0.60 |
| 71 | Plant height | 0.26 | 0.4 | -0.06 | -0.63 |
| | Leaf area index | 0.19 | 0.22 | -0.11 | -0.18 |
| | NDVI | 0.21 | 0.07 | -0.22 | -0.48 |
| | GNDVI | 0.10 | 0.21 | 0.18 | 0.68 |
| | TGI | 0.64 | 0.81 | -0.38 | -0.61 |
| | NDRE | 0.21 | 0.22 | 0.19 | 0.64 |
| | REIP | 0.28 | 0.07 | 0.25 | 0.99 |
| 80 | Plant height | 0.45 | 0.40 | -0.24 | -0.64 |
| | Leaf area index | 0.34 | 0.35 | -0.22 | -0.49 |
| | NDVI | 0.28 | 0.05 | -0.13 | -0.38 |
| | GNDVI | 0.35 | 0.30 | -0.27 | -0.38 |
| | TGI | 0.28 | 0.04 | -0.15 | -0.46 |
| | NDRE | 0.32 | 0.15 | -0.10 | -0.22 |
| | REIP | 0.30 | 0.006 | -0.08 | -0.14 |
| 87 | Plant height | 0.42 | 0.25 | -0.30 | -0.73 |
| | Leaf area index | 0.46 | 0.37 | -0.27 | -0.49 |
| | NDVI | 0.39 | 0.05 | -0.21 | -0.46 |
| | GNDVI | 0.50 | 0.86 | -0.27 | -0.60 |
| | TGI | 0.36 | 0.82 | -0.18 | -0.37 |
| | NDRE | 0.50 | 0.23 | -0.24 | -0.55 |
| | REIP | 0.56 | 0.03 | -0.24 | -0.53 |

Table S5: Mean and standard error (SE) of genomic prediction accuracies obtained from various models and scenarios used for plant height at each time point. The scenario represents the cross-validation scheme used in Figure 1. The $p$-value was derived from a statistical significance test between random regression (RR) and single-time point (ST) GBLUP. Days after sowing in the multi-trait genomic best linear unbiased prediction (MT-GBLUP) model refers to the time point that was coupled with end-of-season plant height.

| Days after sowing | Scenario | Model | Mean | SE | $p$-value |
|---|---|---|---|---|---|
| 45 | CV1 | RR-GBLUP | 0.27 | 0.12 | 0.033 |
| | | ST-GBLUP | 0.33 | 0.12 | |
| 57 | CV1 | RR-GBLUP | 0.32 | 0.17 | 0.080 |
| | | ST-GBLUP | 0.38 | 0.15 | |
| 65 | CV1 | RR-GBLUP | 0.46 | 0.13 | 0.099 |
| | | ST-GBLUP | 0.49 | 0.12 | |
| 71 | CV1 | RR-GBLUP | 0.52 | 0.12 | 0.440 |
| | | ST-GBLUP | 0.53 | 0.11 | |
| 80 | CV1 | RR-GBLUP | 0.65 | 0.11 | 0.360 |
| | | ST-GBLUP | 0.65 | 0.10 | |
| 87 | CV1 | RR-GBLUP | 0.65 | 0.10 | 0.350 |
| | | ST-GBLUP | 0.66 | 0.09 | |
| 71 | CV2 | RR-GLUP | 0.78 | 0.015 | NA |
| 80 | CV2 | RR-GLUP | 0.46 | 0.058 | NA |
| 87 | CV2 | RR-GLUP | 0.17 | 0.050 | NA |
| 80 | CV2.1 | RR-GLUP | 0.84 | 0.023 | NA |
| 87 | CV2.1 | RR-GLUP | 0.73 | 0.031 | NA |
| 87 | CV2.2 | RR-GLUP | 0.86 | 0.010 | NA |
| 45 | MT-CV3 | MT-GBLUP | 0.17 | 0.050 | NA |
| 57 | MT-CV3 | MT-GBLUP | 0.64 | 0.020 | NA |
| 65 | MT-CV3 | MT-GBLUP | 0.76 | 0.009 | NA |
| 71 | MT-CV3 | MT-GBLUP | 0.82 | 0.007 | NA |
| 80 | MT-CV3 | MT-GBLUP | 0.90 | 0.009 | NA |

Table S6: Mean and standard error (SE) of genomic prediction accuracies obtained from multi-trait genomic best linear unbiased prediction (GBLUP) for the different scenarios. DAS and Trait represent days after sowing and the secondary trait coupled with seed yield, respectively. The $p$-value is the significance obtained from the corrected resampled t-test comparing the multi-trait model with the single trait GBLUP on seed yield.

| **DAS** | **Trait** | **CV1** | | | **CV2** | | |
|---|---|---|---|---|---|---|---|
| | | Mean | SD | $p$-value | Mean | SE | $p$-value |
| 45 | Plant height | 0.44 | 0.12 | 0.17 | 0.41 | 0.12 | 0.40 |
| | Leaf area index | 0.42 | 0.11 | 0.41 | 0.41 | 0.11 | 0.23 |
| | NDVI | 0.40 | 0.11 | 0.20 | 0.41 | 0.11 | 0.32 |
| | GNDVI | 0.42 | 0.11 | 0.47 | 0.41 | 0.11 | 0.36 |
| | TGI | 0.39 | 0.11 | 0.19 | 0.41 | 0.11 | 0.37 |
| | NDRE | 0.43 | 0.12 | 0.37 | 0.41 | 0.10 | 0.44 |
| | REIP | 0.43 | 0.11 | 0.35 | 0.41 | 0.10 | 0.41 |
| 57 | Plant height | 0.35 | 0.14 | 0.03 | 0.43 | 0.12 | 0.31 |
| | Leaf area index | 0.42 | 0.11 | 0.45 | 0.41 | 0.11 | 0.11 |
| | NDVI | 0.37 | 0.12 | 0.10 | 0.40 | 0.12 | 0.18 |
| | GNDVI | 0.44 | 0.12 | 0.27 | 0.41 | 0.11 | 0.42 |
| | TGI | 0.43 | 0.13 | 0.40 | 0.41 | 0.12 | 0.38 |
| | NDRE | 0.44 | 0.12 | 0.27 | 0.41 | 0.11 | 0.40 |
| | REIP | 0.44 | 0.12 | 0.27 | 0.41 | 0.11 | 0.44 |
| 65 | Plant height | 0.34 | 0.14 | 0.01 | 0.43 | 0.12 | 0.32 |
| | Leaf area index | 0.40 | 0.12 | 0.16 | 0.40 | 0.11 | 0.13 |
| | NDVI | 0.40 | 0.12 | 0.33 | 0.39 | 0.12 | 0.17 |
| | GNDVI | 0.44 | 0.12 | 0.33 | 0.42 | 0.11 | 0.49 |
| | TGI | 0.43 | 0.12 | 0.36 | 0.40 | 0.13 | 0.35 |
| | NDRE | 0.46 | 0.13 | 0.18 | 0.42 | 0.11 | 0.48 |
| | REIP | 0.44 | 0.13 | 0.30 | 0.41 | 0.11 | 0.45 |
| 71 | Plant height | 0.34 | 0.15 | 0.04 | 0.43 | 0.12 | 0.32 |
| | Leaf area index | 0.40 | 0.11 | 0.11 | 0.40 | 0.11 | 0.11 |
| | NDVI | 0.38 | 0.13 | 0.25 | 0.40 | 0.13 | 0.31 |
| | GNDVI | 0.42 | 0.11 | 0.44 | 0.41 | 0.11 | 0.30 |
| | TGI | 0.41 | 0.13 | 0.48 | 0.41 | 0.12 | 0.44 |
| | NDRE | 0.42 | 0.12 | 0.48 | 0.42 | 0.10 | 0.47 |
| | REIP | 0.41 | 0.13 | 0.48 | 0.43 | 0.10 | 0.33 |
| 80 | Plant height | 0.37 | 0.14 | 0.17 | 0.42 | 0.12 | 0.38 |
| | Leaf area index | 0.39 | 0.13 | 0.24 | 0.40 | 0.12 | 0.23 |
| | NDVI | 0.37 | 0.13 | 0.08 | 0.39 | 0.12 | 0.20 |
| | GNDVI | 0.37 | 0.13 | 0.11 | 0.39 | 0.12 | 0.16 |
| | TGI | 0.37 | 0.13 | 0.09 | 0.39 | 0.12 | 0.21 |
| | NDRE | 0.38 | 0.12 | 0.08 | 0.40 | 0.12 | 0.16 |
| | REIP | 0.39 | 0.12 | 0.12 | 0.40 | 0.12 | 0.16 |
| 87 | Plant height | 0.40 | 0.15 | 0.35 | 0.44 | 0.11 | 0.17 |
| | Leaf area index | 0.39 | 0.14 | 0.25 | 0.40 | 0.13 | 0.25 |
| | NDVI | 0.37 | 0.13 | 0.10 | 0.40 | 0.12 | 0.23 |
| | GNDVI | 0.37 | 0.15 | 0.17 | 0.40 | 0.13 | 0.27 |
| | TGI | 0.37 | 0.13 | 0.08 | 0.40 | 0.12 | 0.18 |
| | NDRE | 0.37 | 0.14 | 0.16 | 0.40 | 0.13 | 0.24 |
| | REIP | 0.38 | 0.14 | 0.19 | 0.39 | 0.13 | 0.22 |

### Figures

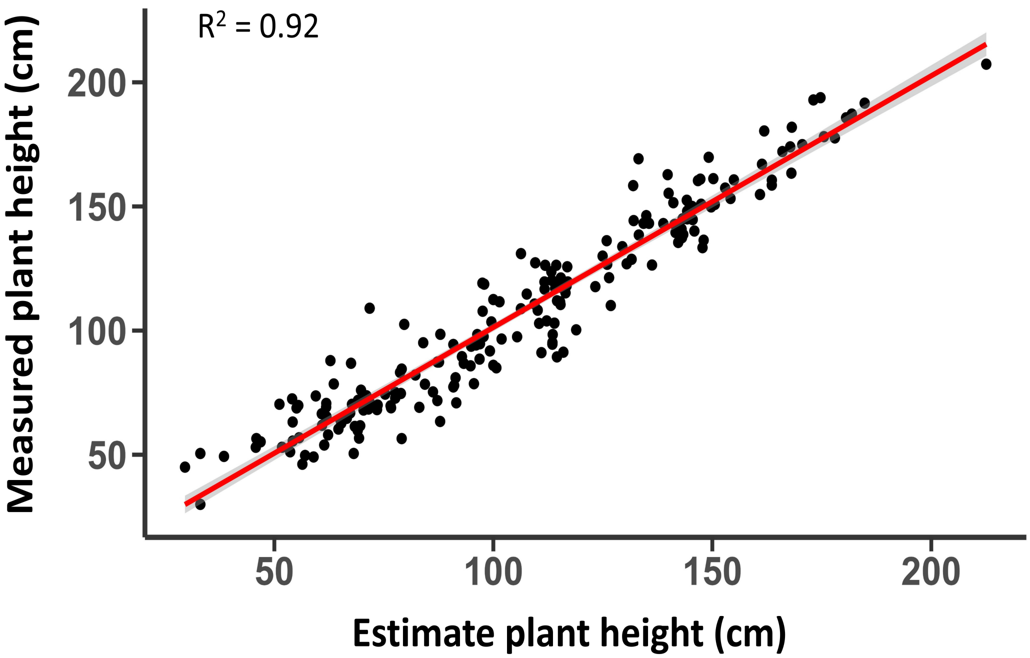


Figure S1: Regression analysis between estimated plant height from image analysis and manually measured plant height in the field.

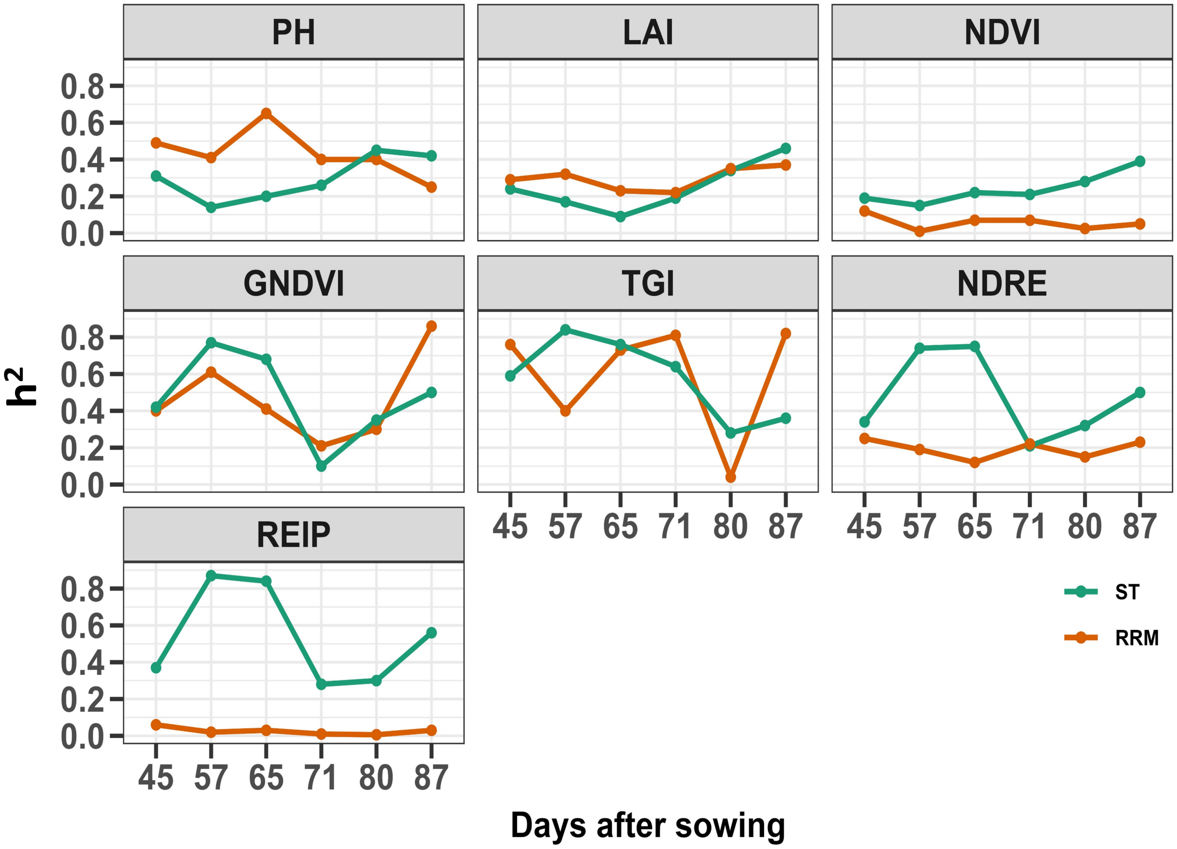


Figure S2: Genomic heritability estimates at each day after sowing for longitudinal traits derived from single time points (green) and random regression (orange) analysis.

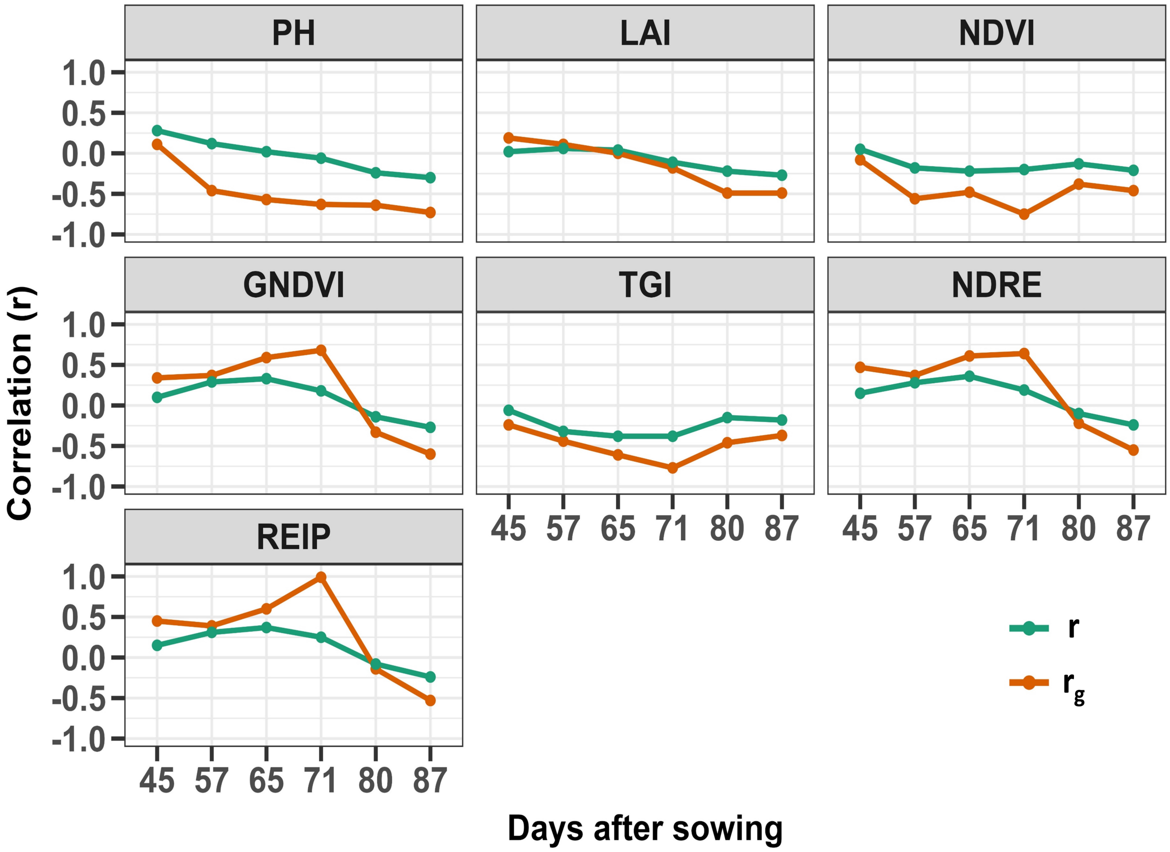


Figure S3: Phenotypic (green) and genetic (orange) correlations between longitudinal traits at each time point and seed-yield.
